# Tau Disaggregation by a CNS-Permeable Small Molecule Reduces Fibril and Oligomer Burden and Preserves Proteostasis and Behavior

**DOI:** 10.64898/2026.06.09.731185

**Authors:** N Mostowfi, R Foreman, J Wang, R Khoury, A Albanese, QL Ma, W Cohn, G Petzinger, M Jakowec, SK Ahmed, PM Seidler

## Abstract

Pathological tau aggregates drive neuronal dysfunction in Alzheimer’s disease (AD) and related tauopathies, yet no approved therapy eliminates existing tau neurofibrillary tangles. Here, we report the development of a coumarin-based small-molecule series that disaggregates tau fibrils and oligomers through a stacking-driven co-assembly mechanism. Structure-activity relationships identified PT-13 as a lead compound that inhibits tau seeding by AD brain-derived matter and reduces aggregate burden measured across both fibrillar and oligomeric tau species. Mechanistic studies demonstrate that disaggregation does not generate soluble oligomeric intermediates, addressing a central question in the field. PT-13 is brain-penetrant and well tolerated in vivo. In a tauopathy mouse model, PT-13 treatment reduces tau pathology while preserving behavioral function, proteasome capacity, and synaptic integrity. These findings establish small-molecule tau disaggregation as a viable therapeutic strategy and provide a molecular framework for the design of aggregate-directed therapeutics in neurodegeneration.

## Introduction

AD is an irreversible neurodegenerative disorder characterized by rampant protein aggregation, encompassing amyloid-β (Aβ) plaques and tau neurofibrillary tangles (NFTs), which impact key brain regions that underlie dementia. Since 2022, three disease-modifying anti-amyloid antibodies have received FDA approval, each of which enhances the removal of pathological Aβ aggregates (1,2). Clinical success in slowing AD progression with these antibodies is the first trace of evidence that targeting and removing pathogenic protein aggregates may alter the trajectory of neurodegenerative disease, underscoring the promise of aggregate-directed therapeutic strategies (2,3). A sole dependence on anti-amyloid therapy, however, has major limitations, since AD is multifactorial, involving aggregation of both Aβ and tau.

Tau is an important unfulfilled drug target, as evidenced by its strong correlation with cognitive decline and brain atrophy (4). Tau NFTs spread autonomously by prion-like seeding mechanisms (5, 6), establishing that tau aggregates themselves are necessary drug targets independent of, or in addition to Aβ plaques and monoclonal antibodies that target them. Additionally, evidence suggests that Aβ could act synergistically, increasing tau aggregation and propagation by seeding (7, 8, 9) further emphasizing the need for tau-targeted therapeutics to manage aggregates that are spawned by amyloid pathology.

A major limitation in drug development for tau and other amyloid proteinopathies has been identifying small molecules that can both disrupt protein aggregates and sufficiently cross the blood-brain-barrier (BBB). Epigallocatechin gallate (EGCG), a polyphenol found in green tea, is a notable in vitro tau inhibitor due to its ability to disaggregate fibrils and thereby prevent prion-like seeding (10). While EGCG lacks brain permeability, deciphering its mechanism has helped uncover druglike compounds that can similarly bind to promote disaggregation.

The Cryo-EM structure of EGCG bound to purified AD tau PHFs revealed how small molecules can bind to protein filaments, leading to their disaggregation (10). A key feature of small molecules that disaggregate tau is their ability to stack as co-filaments, ultimately driving fibril disassembly. Subsequent work showed that protein aggregates can nucleate small-molecule co-assemblies, and that the conformational strain imposed by bound small molecules as they relax into their own filamentous structures upon reaching a critical mass can drive aggregate remodeling and disassembly (11). With this knowledge, thousands of Central Nervous System (CNS) compounds have been screened in their potential to bind to, and form filament co-assemblies with structure-based pharmacophores on tau and other amyloid fibrils.

Here, we introduce a planar, coumarin-based chemical series of small molecules with the ability to stack and disassemble tau aggregates. A key distinction of this chemical series from other small-molecule tau disaggregants is its favorable CNS drug-like properties and blood-brain barrier penetration. Given the promising pharmacological properties and tau disaggregation activity of this series, we investigated its feasibility for in vivo testing in a tauopathy mouse model.

A major outstanding question in the field has been whether small-molecule tau disaggregants produce beneficial therapeutic outcomes or, alternatively, promote fibril disassembly into oligomeric species with greater toxicity. Our results show that the coumarin-based tau disaggregant is well tolerated in tauopathy animals, producing functional benefits accompanied by reductions in tau aggregate species. Collectively, these findings support the hypothesis that aggregate disassembly is a viable therapeutic mechanism in the context of proteinopathies.

## Results

### Determinants of Tau Seeding Inhibition and Fibril Disaggregation

N-(6-methyl-2-pyridinyl)-2-oxo-2H-chromene-3-carboxamide (CNS-16) is a coumarin-based small molecule inhibitor identified by screening using tau seeding assays (10). The coumarin moiety is linked to a 6-methyl pyridine, creating favorable potential stacking interactions, a determinant for fibril binding and disassembly. The coumarin backbone also confers high reported bioavailability and the amide-linked pyridine shares features with certain nitrogen-rich tau PET ligands, such as T807 and MK-6240 (10, 12, 13, 14). We explored the structure-activity relationship of CNS-16, synthesizing 40 coumarin-based analogs using the reaction scheme shown in Fig. 1. Amongst the series, PT-13 emerged as a top interest, inhibiting tau seeding and disaggregating fibrils. PT-13 differs from CNS-16 by a cyano substituent on the pyridine, replacing the methyl substituent.

**Figure 1.**
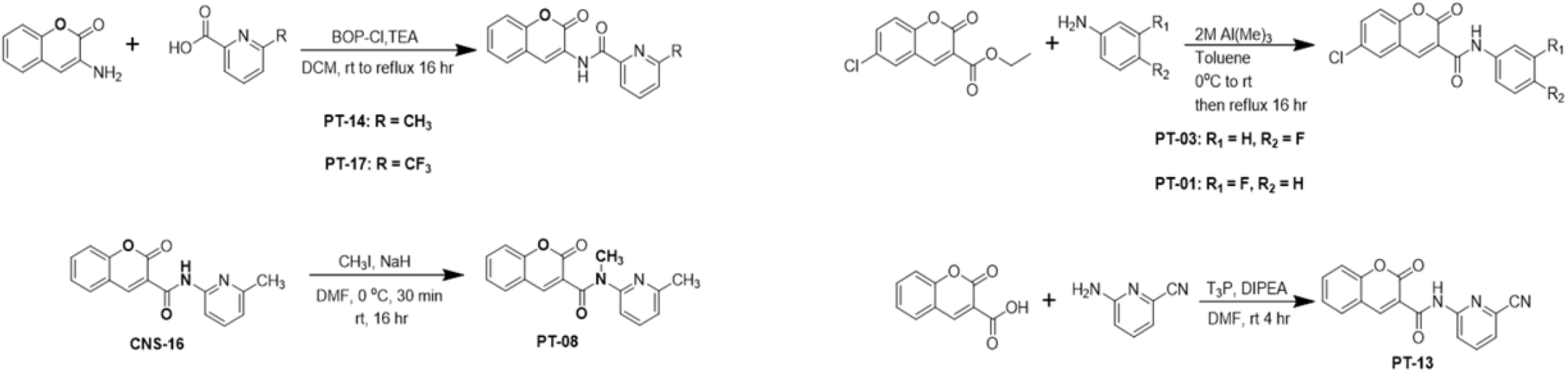
Chemical synthesis scheme for coumarin-based tau disaggregant series. Analogs were synthesized using classic amide coupling schemes to obtain the initial series of compounds derived from CNS-16 to uncover the hit-lead compound, PT-13.

Among the coumarin series of analogs synthesized by amide coupling, PT-13 showed significant activity as a tau inhibitor in seeding assays using K18 biosensor cells (Fig. 2). Tau biosensor cell assays were conducted by treating crude AD brain homogenates with the coumarin-based small molecule series, and residual seeding power by AD tau aggregates was surmised by subsequent transfection with inhibitor-treated brain homogenate seeds. Quantifying puncta from seeded cells revealed that PT-13 inhibits tau seeding with greater efficacy than CNS-16 (Fig. 2 A-D). The nitrile (cyano) substituent on PT-13 is a bioisostere with relatively greater metabolic stability and bioavailability compared to carbonyl groups, carboxylic acids, halogens, and hydroxyls, and has found uses in drug development in leukemia and cardiovascular disease (15,16). Although, the activity of cyano-containing tau inhibitors had not been extensively explored.

**Figure 2.**
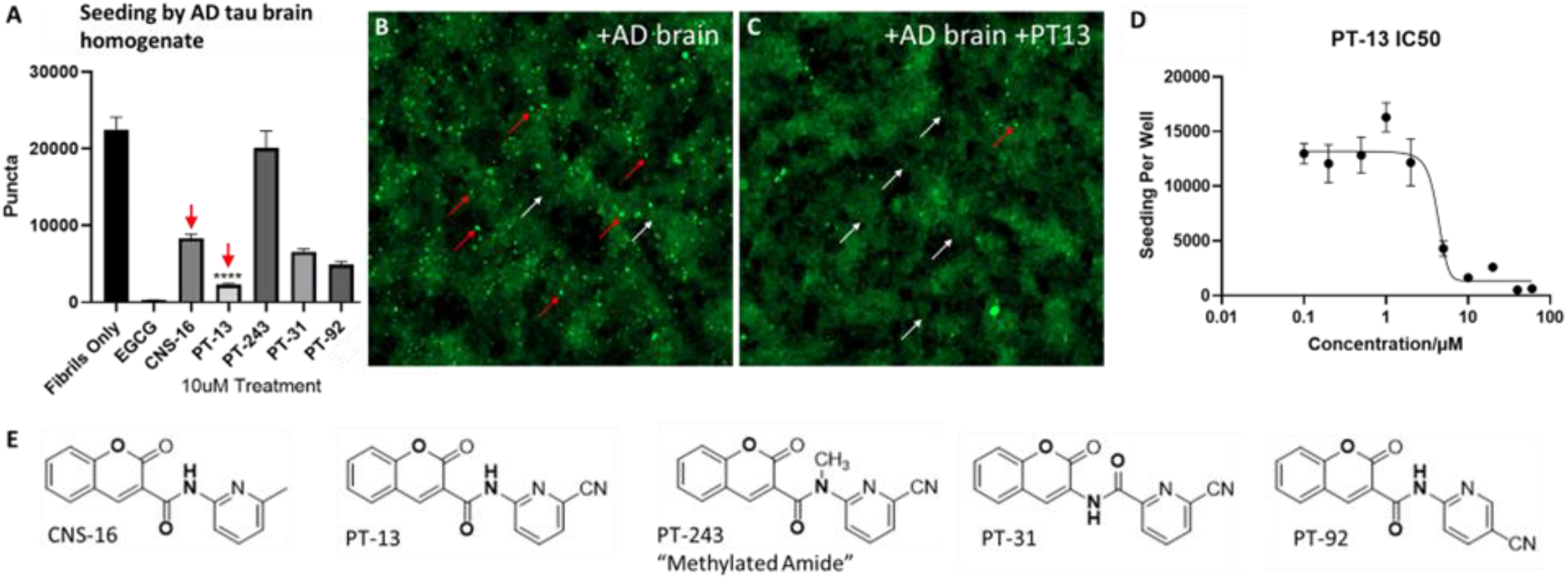
PT-13 is a Tau Seeding Inhibitor. (A) Tau K18 biosensor cell seeding assays. AD crude brain homogenate was used as a seed following overnight incubation with coumarin-based molecules, as indicated. The number of puncta were normalized to cell confluence and quantified to assess seeding inhibition. Each condition contains a final concentration of 10 µM coumarin-based molecule on cells. EGCG was included as a reference tau inhibitor molecule. (B and C) Images from seeded cells without (B) or with (C) inhibitor. (D) Dose titration with PT-13 inhibits seeding with low micromolar potency. (E) Chemical structures of CNS-16 analogs. (*\*\*\*\* indicates p < 0*.*0001*; n = 3 per sample).

Supplementary Table 1 reports the measured tau inhibitor seeding activity for the full series of 40 coumarin-based tau inhibitors synthesized and tested. Chemical structures and corresponding seeding inhibition data for a small subset of cyano-pyridine analogs are shown in Fig. 2A and E. Compounds with activities approaching PT-13 included PT-31, a reverse-amide analog of PT-13, and PT-92, which contains a cyanopyridine substituent at the para position. Crucially, PT-243, an analog of PT-13 with an N-methyl amide that disrupts intermolecular stacking, displays no tau inhibitor activity. These findings indicate that both the cyano substituent and intermolecular stacking interactions are key determinants of the enhanced tau inhibitory activity of PT-13.

Given the sensitivity of tau inhibitory activity to chemical modifications that disrupt stacking interactions, we assessed the fibril disaggregation activity of PT-13 and PT-243 by quantitative electron microscopy (qEM). As shown in Fig. 3A and B, incubating AD brain-purified tau fibrils with PT-13 resulted in a sixfold reduction in fibril density relative to fibrils treated with PT-243, which exhibited no change in density. Representative negative-stain EM images shown in Fig. 3C and D reveal that the limited fibrils remaining after PT-13 treatment appear qualitatively shorter than untreated fibrils. These data are congruent with tau seeding assays in Fig. 2, which show reduced seeding by PT-13 but not PT-243. Collectively, these findings indicate that both fibril disaggregation and seeding inhibition are disrupted by the N-methyl amide substituent, which impedes intermolecular stacking.

**Figure 3.**
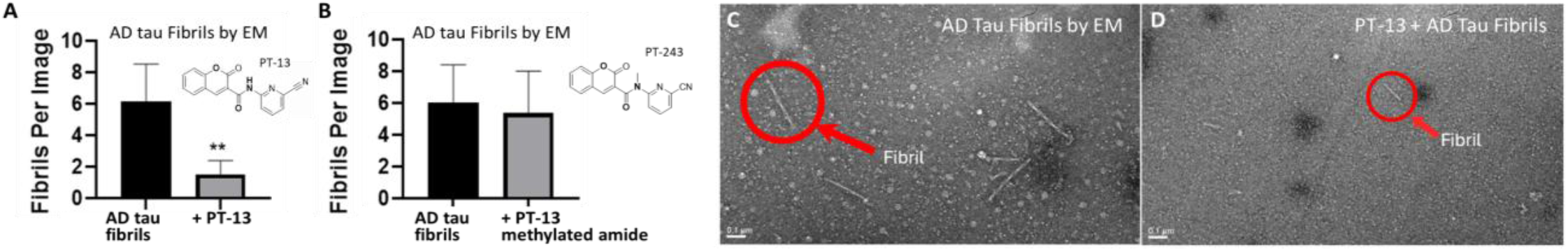
Fibril Disaggregation Activity Tracks with Drug Stacking. (A and B) AD brain-purified tau fibrils were treated with PT-13, or the N-methyl amide analog, PT-243. AD tau filaments were counted from 25 negatively-stained electron micrographs to quantify remaining fibril density as a function of inhibitor treatment. AD purified fibrils incubated with PT-13 compared to fibrils alone led to a significant reduction in fibrils per image, A, compared to fibrils treated with PT-243, in B. (C and D) Representative negative-stained electron micrographs. Tau fibrils that remain after PT-13 treatment, in D, are qualitatively shorter compared to non-treated fibrils in C. (*\*\*\*\* indicates p < 0*.*0001*; n = 25 images for qEM).

### PT-13 Decomposes Tau Oligomers and Inhibits Associated Seeding

It is conceivable that small molecules that disaggregate tau fibrils could generate soluble tau oligomers capable of exerting proteopathic effects (17). To assess the sensitivity of tau oligomers to PT-13, we probed AD brain homogenate immunoreactivity using the tau oligomer-selective monoclonal antibodies TTC-M2, TOMA-1, TOMA-2, and TOMA-3, which recognize a diverse range of tau oligomer species (18, 19). Immunoblotting data shown in Fig. 4 demonstrate that tau oligomer immunoreactivity undergoes a dose-dependent reduction with increasing PT-13 concentration. These data confirm that disaggregant treatment does not increase oligomer formation across a wide range of PT-13 concentrations and instead demonstrate that tau oligomers, like fibrils, can be decomposed by the same small-molecule disaggregants.

**Figure 4.**
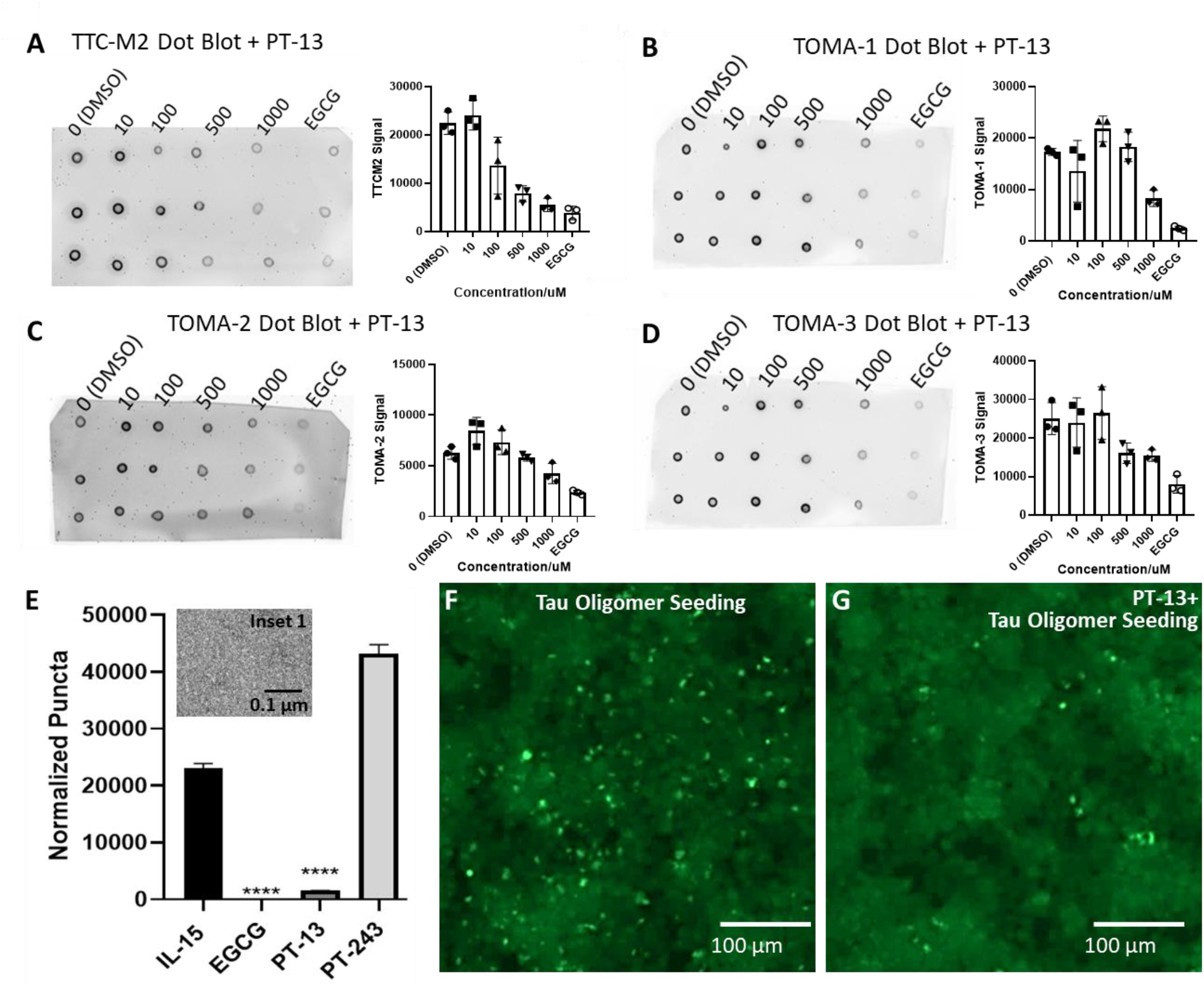
PT-13 Disaggregates Tau Oligomer Species. (A-D) Crude AD brain homogenate treated with increasing PT-13 dose reduces immunoreactivity to a range of tau oligomer antibodies. Dot blots with triplicate measures are shown alongside corresponding densitometry plots. (E-G) Seeding by tau K18+ recombinant oligomer and oligomer seeding inhibition by PT-13 treatment (representative image shown in G). (\*\*\*\**p* < 0.0001; n = 3 per condition). Inset 1 shows negative-stain EM of IL15 oligomer.

To characterize the extent to which PT-13-mediated disaggregation affects oligomer-induced seeding, we generated recombinant tau oligomers using Ionic Liquid 15 (IL15; 1-n-butyl-3-methylimidazolium n-octylsulfate), a previously established model for generating seeding-competent tau oligomers (20). As shown in Fig. 4E and F, tau-K18+ oligomers strongly seed tau biosensor cells despite an evident lack of observable fibrils assessed by negative-stain EM (Fig. 4E, Inset 1). PT-13 treatment dramatically reduced seeding by recombinant tau oligomer, while PT-243 had no effect. These results align with immunoblotting data demonstrating that PT-13 disrupts oligomeric tau species, indicating that small-molecule disaggregants can effectively dismantle oligomer structures and suppress their seeding activity.

### PT-13 CNS Druglike Characteristics

We assessed the CNS druglike properties of PT-13 by in vitro ADMET properties and pilot studies. As shown by the data table in Supplementary Figure 1A, PT-13 displays permeability in MDCK-MDR1 cells, an in vitro model for BBB permeability. MDCK-MDR1 cells express P-glycoprotein (P-gp), which is involved in drug efflux. PT-13 displayed an experimental efflux ratio of 1.0 in MDCK-MDR1 permeability assays indicating effective membrane crossing and retention in cells. Consistent with this data, PT-13 has a measured LogD of 2.7, indicating excellent cell permeability. On this basis, we conducted pilot studies investigating BBB permeability. To determine the superior delivery method, C57BL/6J mice were injected with the highest tolerated dose of PT-13 at 15 mg/kg subcutaneously (SC) or 5 mg/kg intraperitoneally (IP) (Supplemental Figure 1B). Quantification of PT-13 concentrations using liquid chromatography-tandem mass spectrometry (LC-MS/MS) determined SC injection to yield superior brain levels, achieving a maximal concentration (Cmax) of over 121.8 ng/g (417 nM). No adverse effects were seen due to PT-13 injection over a 29-day period (Supplementary Figure 1 D-H).

### PT-13 Treatment Effects

The promising initial in vivo results prompted expanded testing to assess the therapeutic effects of PT-13 treatment in a tauopathy mouse model. These studies utilized the rTg4510 tauopathy mouse model, in which tau pathology first emerges in the hippocampus and entorhinal cortex prior to spreading to neocortical regions (21, 22). The formation and pattern resemble tau pathology in AD making rTg4510 mice a useful proxy for evaluating the effects of tau disaggregants in vivo. rTg4510 mice received PT-13 or vehicle by SC injection twice daily due to a short PT-13 half-life of less than 2 hrs (Supplementary Figure 1 C). Injections were made three days per week, beginning at three months of age, as tau pathology accelerates markedly between four and six months of age in this model (23).

Behavioral studies were conducted at three months of age to establish baseline performance and at six months of age to assess the functional impact of PT-13 treatment. As shown in Fig. 5A and B, PT-13 treatment improved performance in cognitive measures assessed using the Y-maze. No differences in Y-maze performance were observed at baseline. Following three months of treatment, PT-13-treated mice exhibited a substantial increase in spontaneous alternation compared to vehicle-treated controls, suggesting preservation of spatial memory, deficits of which emerge in rTg4510 mice beginning around four months of age (35). No differences were detected in Morris Water Maze or Probe Test (Supplementary Fig. 2). We also assessed motor impairment using the rotarod performance assay (Fig. 5). At six months of age, PT-13-treated mice exhibited increased latency to fall relative to vehicle-treated controls, indicating improved motor performance. Collectively, these functional readouts suggest that disaggregant treatment is well tolerated and provides benefits to tauopathy mice across a range of cognitive and motor domains.

**Figure 5.**
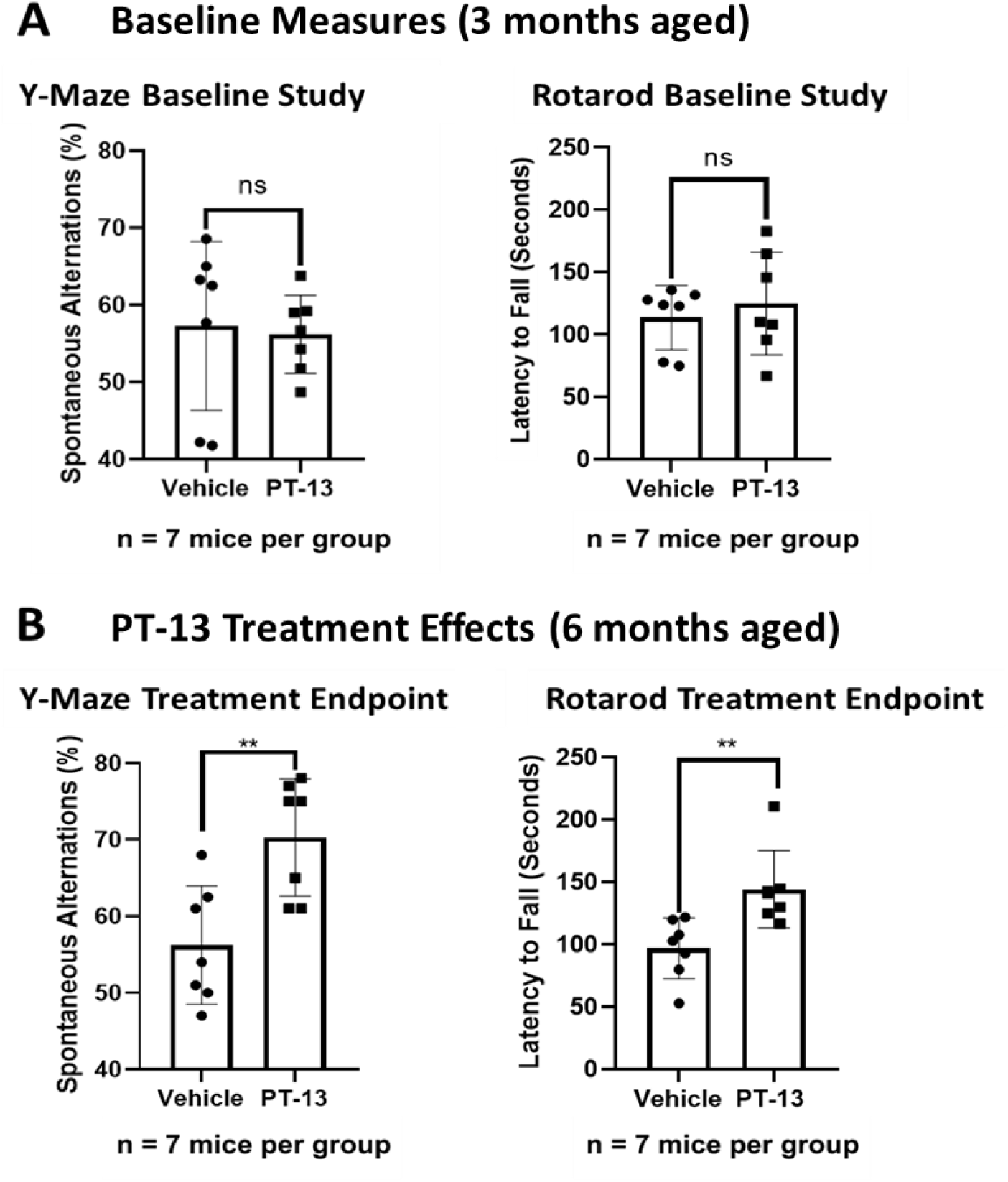
PT-13 treatment protects against functional decline in the rTg4510 tauopathy mouse model. (A) Baseline behavioral assessments performed at 3 months of age showed no significant differences between treatment groups in the Y-maze and rotarod assays (A-C). (B) After three months of PT-13 treatment, at 6-months aged, PT-13-treated mice exhibited increased spontaneous alternation in the Y-maze, indicative of improved spatial memory (p < 0.005; n = 7 mice per group). In accelerating rotarod experiments, PT-13-treated mice had a greater latency to fall compared to vehicle-treated controls (p < 0.005; n = 7 mice per group).

### Neuropathological Effects of PT-13 treatment

Following the observed improvements in memory and motor function, the treatment study was terminated to assess corresponding neuropathological outcomes. Immunohistochemical analysis revealed a trend toward reduced AT8 immunoreactivity in the hippocampal CA1 region following PT-13 treatment (Fig. 6 A-B). In addition, immunoreactivity assessed using a commercial TOMA antibody (Sigma-Aldrich, Catalog # MABN819) reduced oligomeric tau in soluble fractions isolated from the cortex and striatum of PT-13-treated animals relative to vehicle controls (Fig. 6 C-D). Tau oligomers are thought to mediate neuronal dysfunction in early tauopathy and AD. The reduction in oligomeric tau, together with the trend towards reduced AT8-positive pathology in the CA1 region could contribute to the improved functional performance following PT-13 treatment.

**Figure 6.**
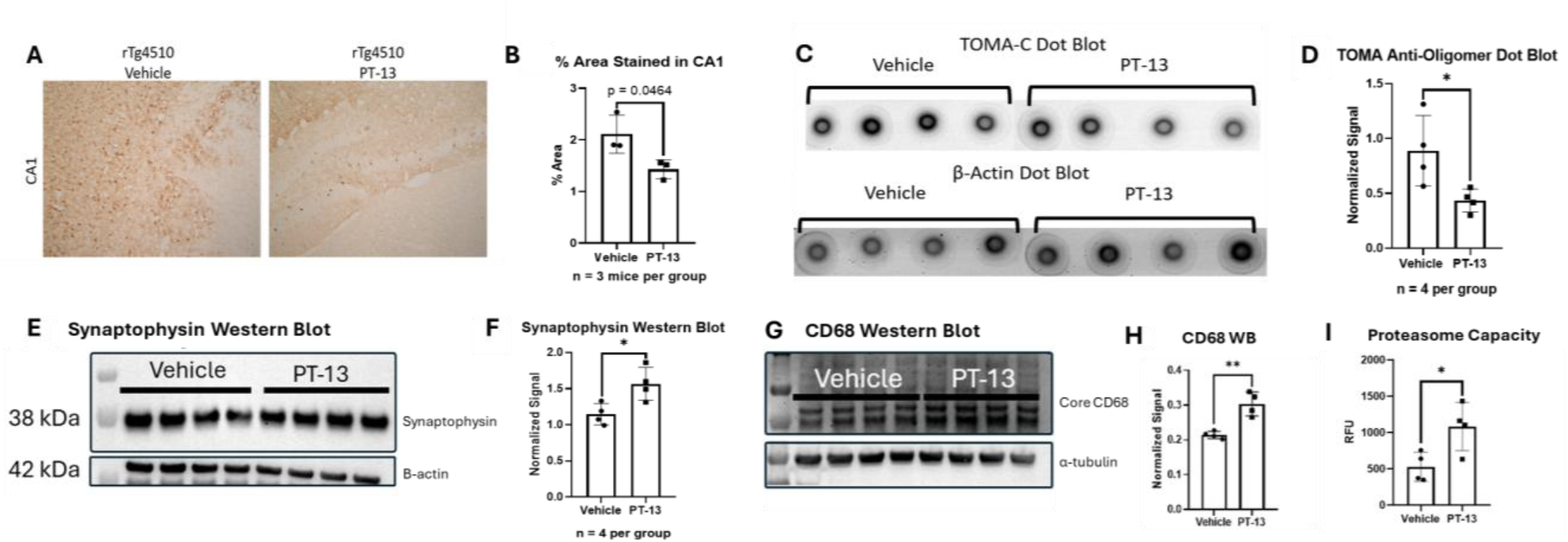
Disaggregant-Mediated Reductions in Tau Aggregate Load and Neuropathologic Markers. (A and B) AT8 IHC of representative hippocampal slices in the CA1 region from PT-13 and vehicle-treated rTg4510 mice. (C and D) Dot blot analysis of oligomeric tau detected using TOMA-C antibody. Representative dot blots show TOMA-C immunoreactivity in the lysis soluble fraction processed from the cortex and striatum from vehicle-treated and PT-13 treated rTg4510 mice normalized to β-Actin. Quantification of normalized TOMA-C signal intensity is shown. PT-13 treatment resulted in a significant reduction in oligomeric tau levels compared to vehicle controls (For IHC: n = 3 mice per group, 2 sections per mouse. For dot blots: *p* < 0.05; *n* = 4 mice per group). (E and F) Western Blot for synaptophysin from cortex tissue of rTg4510 mice and corresponding quantification (*p* < 0.05; *n* = 4 mice per group). (G and H) Western Blot for CD68, a marker for microglial phagocytosis activity and corresponding quantification (*p* < 0.0025; *n* = 4 mice per group). (I) Proteasome activity from rTg4510 brain homogenate measured by fluorescence intensity using Suc-LLVY-AMC for vehicle-treated and PT-13 treated mice.

These findings provide evidence that PT-13 promotes tau disaggregation in vivo without evidence of increased toxicity.

Expanding the scope of our study, we assessed synaptic protein markers as a functional index of synaptic integrity and PT-13 treatment. Tau pathology is known to induce synaptic deficits, including reductions in presynaptic vesicle mobility and impairment of postsynaptic dendritic spines (26, 27). Presynaptic integrity was assessed by synaptophysin western blotting. Synaptophysin is an abundant presynaptic vesicle protein commonly used as a marker of presynaptic terminals and synaptic integrity. PT-13-treated animals exhibited increased synaptophysin levels in cortical lysates relative to vehicle-treated controls (Fig. 6E-F), suggesting that tau disaggregation preserves presynaptic integrity and protects against tau-mediated synaptic dysfunction.

Postsynaptic integrity was assessed by western blotting for postsynaptic density protein 95 (PSD-95), a scaffolding protein critical for recruiting and anchoring NMDA and AMPA receptors at the postsynaptic membrane (28, 29). Western blot analysis shown in Supplementary Figure 3 revealed a trend toward increased PSD-95 in the SDS-insoluble fraction and reduced PSD-95 in the RIPA buffer soluble fraction from cortical tissue of PT-13-treated mice. Since PSD-95 functions as a core scaffolding protein within the postsynaptic density, this shift may reflect increased incorporation of PSD-95 into mature postsynaptic complexes. On this basis, it is possible that reduction of tau aggregates by PT-13 alleviates synaptic dysfunction, allowing for the formation of postsynaptic structures and recruitment of PSD-95 to receptor-containing complexes.

Tau aggregate clearance is mediated by multiple proteostatic pathways, including microglial phagocytosis (30) and proteasome-dependent degradation (31). We therefore assessed markers of protein clearance, since disaggregant treatment could help preserve proteostatic functions. Prior studies show that as tau pathology progresses, sustained microglial activation can drive a transition towards a dysfunctional, hyporeactive state characterized by impaired phagocytic capacity, reduced aggregate clearance, and further accumulation of tau oligomers and fibrils (30, 32). Similarly, proteasome activity declines in AD and related proteinopathies, contributing to impaired elimination of pathogenic protein aggregates (33).

To assess microglial phagocytic capacity, we measured CD68 levels by western blotting. CD68 is a lysosomal and phagocytic marker associated with activated microglia. PT-13 treatment produced a statistically significant increase in CD68 levels in cortical lysates relative to vehicle-treated rTg4510 mice (Fig. 6 G-H). To assess proteasome capacity, we measured proteasome activity in enriched 26S proteasome fractions isolated from PT-13- and vehicle-treated rTg4510 brain homogenates using the fluorogenic substrate Suc-LLVY-AMC. As shown in Fig. 6I, PT-13-treated mice exhibited increased proteasome activity relative to vehicle-treated controls.

Collectively, in vivo treatment shows that PT-13 protects rTg4510 mice, preserving behavioral function across cognitive and motor domains. Neuropathological measures of tau pathology were similarly improved, with reduced oligomer load and decreased AT8 immunostaining. These benefits extend to cellular outcomes, including preservation of synaptic integrity, improved microglial function, and enhanced proteasome activity, supporting the feasibility of tau disaggregation as a therapeutic strategy.

## Discussion

Current treatments for AD are primarily focused on reducing Aβ pathology, and no FDA-approved therapies directly induce tau disaggregation to eliminate existing neurofibrillary tangles (NFTs). Although Aβ-directed therapies such as lecanemab and donanemab provide clinical benefit, they do not entirely halt disease progression, underscoring the need for therapeutic approaches that directly target tau pathology (1,3,34). Existing small molecules targeting tau largely function by inhibiting aggregation, attempting to prevent the conversion of monomeric tau into fibrillar species. However, these approaches do not remove pre-existing NFTs (1, 35, 36). Persistent and/or pre-existing tau aggregates may continue to drive neuroinflammatory responses, including microglial dysfunction and impaired proteasome function, thereby contributing to ongoing neuronal stress (30, 37). The ability to eliminate existing NFTs is therefore a critical feature of an effective tau-directed therapeutic strategy. This rationale motivated our development of a small-molecule disaggregase approach.

Here, we show that PT-13, a member of a coumarin-based ligand series, potently inhibits tau seeding through disaggregation of both fibrillar and oligomeric tau species. The favorable CNS drug-like properties and brain permeability of PT-13 enabled in vivo testing, which revealed benefits spanning functional, cellular, and neuropathological domains. These findings are particularly important because a longstanding concern surrounding aggregate disassembly has been the possibility that fibril disruption could generate soluble oligomeric species with enhanced toxicity. In contrast, our data demonstrates that PT-13 not only disaggregates fibrils, but also reduces oligomer immunoreactivity and oligomer-induced seeding. Consistent with these observations, PT-13 treatment produced no evidence of increased oligomer burden in vitro or in vivo. Together, these findings suggest that tau disaggregation can be achieved without exacerbating proteinopathy and support aggregate disassembly as a viable therapeutic mechanism.

The coumarin series further provides insight into the molecular features that drive disaggregation by small molecules. The addition of a nitrile substituent on the amide-linked pyridine emerged as a key determinant of activity and may functionally substitute for aromatic hydroxyl groups that contribute to disaggregation by polyphenols such as EGCG (10). Another critical determinant is intermolecular stacking, which appears necessary for fibril binding and disaggregation. Chemical modification specifically designed to disrupt intermolecular stacking among PT-13 molecules abolished both fibril disaggregation and seeding inhibition. These findings provide mechanistic insight into small-molecule-mediated aggregate disassembly and establish design principles for future generations of tau disaggregants.

Beyond reducing aggregate burden, PT-13 treatment was associated with preservation of presynaptic integrity, increased markers of microglial phagocytic capacity, and enhanced proteasome activity in tauopathy mice. Since both microglial clearance and proteasome-mediated degradation are compromised during disease progression, these findings support the hypothesis that tau disaggregation may help restore proteostatic pathways that become impaired during proteinopathy.

Overall, these findings establish PT-13 as a brain-penetrant small-molecule tau disaggregase that produces beneficial outcomes in a tauopathy model while avoiding increased oligomer loads or proteotoxicity. These studies broadly support the concept that therapeutic disassembly of pathogenic protein aggregates represents a feasible strategy for treating tauopathies and related neurodegenerative disorders.

## Materials and Methods

### Chemistry

Molecular properties for the initial series of analogs for hit-lead SAR were calculated using StarDrop software (Optibrium Ltd., Cambridge, UK, version 8). The initial CNS-16 and subsequent PT-13 analogs were synthesized as described in our international patent (38).

### Preparation of AD crude brain extracts and purified AD tau fibrils

Human autopsy samples from the University of California, Los Angeles (UCLA) Pathology Department were obtained according to US Department of Health and Human Services regulations from patients consenting to autopsy. Following established protocols (10, 39), sections of approximately 250 mg of autopsy brain tissue from human AD patients were cut and homogenized using a Polytron in 750 μL of sucrose buffer (0.8 M NaCl, 10% sucrose, 10 mM Tris-HCl, pH 7.4) supplemented with 1 mM EGTA and 5 mM EDTA. Homogenates, flash-frozen and stored at −80°C, were used directly as tau seeds in biosensor cell assays. To obtain purified tau fibrils from crude brain homogenates, sections of approximately 1 g of autopsy brain tissue from human AD patients were cut and homogenized using a Polytron in 15 mL of sucrose buffer (0.8 M NaCl, 10% sucrose, 10 mM Tris-HCl, pH 7.4) supplemented with 1 mM EGTA and 5 mM EDTA. Homogenates were centrifuged at 20,100g for 20 min at 4°C, and the supernatants were collected and transferred to airfuge ultra-tubes and spun at 95,000 rpm for 60 min. The ultra-pellet was resuspended in 2.5 mL of sucrose buffer supplemented with 5 mM EDTA and 1 mM EGTA and centrifuged at 20,100g for 30 min at 4°C. The supernatant was spun at 95,000 rpm for 60 min at 4°C, and the pellet containing fibrils was resuspended in 100 μL of 20 mM Tris-HCl (pH 7.4) and 100 mM NaCl.

### HEK293T biosensor seeding assays

Cell seeding assays were performed using HEK293T cells stably expressing the K18 aggregation-prone tau fragment containing residues 244 to 372 with two point mutations (P301S and V337M) fused at the C-terminus with green fluorescent protein (GFP) (39). Cells were cultured in Dulbecco’s Modified Eagle’s Medium (DMEM) (Life Technologies, catalog no. 11965092) supplemented with 10% (v/v) fetal bovine serum (FBS) (Life Technologies, catalog no. A3160401), 1% penicillin/streptomycin (Life Technologies, catalog no. 15140122), and 1% GlutaMAX (Life Technologies, catalog no. 35050061) in a T25 flask at 37°C in a humidified incubator with 5% CO_2_.

### Tau biosensor dose titrations

K18-CY cells were plated in a 96-well dish in a volume of 100 μL 24 hours before transfection. Compounds to be tested were preincubated with crude AD brain homogenate [1 μL of crude brain homogenate diluted with 19 μL of Opti-MEM (Thermo Fisher Scientific, catalog no. 31985062)] at 4°C overnight before transfecting onto cells grown to 60-80% confluence. For the dose-response assay, twelve concentrations were selected: 0, 0.1, 0.2, 0.5, 1, 2, 5, 10, 20, 40, 60, and 80 μM, and each condition was carried out in triplicate. To enhance seeding, the crude brain homogenates were sonicated in a Qsonica multiple-horn water bath for 3 min at 40% power before transfection. For transfection, Lipofectamine 2000 (Thermo Fisher Scientific, catalog no. 11668019) was used according to the manufacturer’s instructions at a 1:20 ratio. Non-transfected cells served as the blank group, and transfected brain homogenate alone served as the positive control for seeding. A BioTek Cytation 5 microscope was used to obtain fluorescent images from seeded 96-well plates. The GFP channel was used, and a 3 × 2 montage mode was set up for the seeded area to capture as much area as possible in each well. Acquired images were analyzed by stitching, preprocessing, and deconvolution for high-content imaging.

### Image analysis

Images were analyzed using ImageJ software to determine the number of tau puncta in each well (39). Background fluorescence from unseeded cells was subtracted, and aggregates were counted as peaks with fluorescence above background using the built-in Particle Analyzer as described previously. Results were normalized to confluence as determined by image analysis. For dose titrations, each concentration was repeated in triplicate and the average number of puncta was calculated from triplicate measures. Statistical analyses were performed using GraphPad Prism software.

### Preparation of negative-stained grids

Purified AD tau fibrils were incubated with 1× phosphate-buffered saline (PBS) or 250 µM of ligand overnight. Six µL of sample was pipetted onto a carbon-coated copper grid (TED PELLA, catalog no. 01754-F) and allowed to incubate for 3 min. Excess liquid was removed by blotting the edge of the grid with filter paper. The grid was then treated with 10 μL of negative-stain solution (4% uranyl acetate) for 2 min before excess stain was removed using filter paper. The grid was air-dried for 10 min before being stored in a grid box for further analysis.

### Acquisition of qEM images

To obtain qEM images, negative-stain EM grids of each sample were screened at a magnification of ×12,000, and images were collected in 5-μm increments. Fibrils were counted from collections of 50 micrographs for each experimental condition, including purified fibrils with and without 250 μM inhibitors incubated at 4°C overnight.

### Dot blot analysis

For dot blots, 2 µL of each crude brain homogenate sample was pipetted onto a nitrocellulose membrane and allowed to dry for 5 minutes before being blocked in 5% bovine serum albumin (BSA) in 1× Tris-buffered saline with 1% Tween 20 (TBST) for 1 hour at room temperature. Membranes were incubated overnight at 4°C with the following primary antibodies: TOMA (Sigma-Aldrich, catalog no. MABN819, 1:1000), T22 (Sigma-Aldrich, catalog no. ABN454, 1:1000), and M204 (1:1000, gifted by Dr. David Eisenberg’s laboratory). Additional primary antibodies (TOMA-1, TOMA-2, TOMA-3, and TTCM2, all at 1:1000) were gifted by Dr. Reza Kayed’s laboratory. After washing three times with TBST, membranes were incubated for 1 hour at room temperature with secondary anti-mouse IgG or secondary anti-rabbit IgG (both Bio-Rad Laboratories, 1:3000) diluted in 5% BSA in TBST. Blots were developed using an enhanced chemiluminescence (ECL) system (Bio-Rad Laboratories, Hercules, CA) and visualized using an iBright imaging device (Thermo Fisher, Waltham, MA).

### Animal Experiments

All animal experiments were approved by the USC Institutional Animal Care and Use Committee and performed under their supervision. For brain permeability and dosage tolerance studies, C57BL/6J mice (Jackson Laboratories: JAX:000664) were used. For the treatment study, rTg4510 mice hemizygous for Tg(Camk2a-tTA)1Mmay and hemizygous for Fgf14<Tg(tetO-MAPT*P301L)4510Kha> (Jackson Laboratories: JAX:024854) were housed on a 12-hour light-dark schedule.

### Dosage Tolerance Study

Pilot study assessing brain permeability and dosage tolerance: C57BL/6J mice received subcutaneous injections of the highest tolerated concentration of PT-13 or vehicle three times per week for 21 days. A separate cohort of uninjected mice was included to confirm the absence of vehicle-related toxicity. To assess potential off-target effects, body weight, food intake, and water consumption were monitored across all groups throughout the dosing period.

Mice were injected subcutaneously (SC) with 15 mg/kg of PT-13 or intraperitoneally (IP) at 5 mg/kg and euthanized by perfusion at 30, 60, and 150 minutes post-injection. Brains were collected by standard dissection and immediately frozen.

### Pharmacokinetics

Analysis of PT-13 brain concentrations was performed at the USC Alfred E. Mann Multi-Omics Mass Spectrometry Core (Director: Whitaker Cohn, Ph.D.). Samples were stored at −80°C and thawed on ice prior to analysis. Drug-naïve brain lysate was spiked with freshly prepared stock solutions of PT-13 at various concentrations (0, 1, 10, 100, and 1000 pmol/tube) to create a five-point standard curve. An internal standard (IS) solution of PT-52 (10 µL, 100 pmol/µL) was added to each sample for quality control to normalize for sample loss. Brain tissue (~100 mg) was homogenized in 4 volumes of ice-cold 80% acetonitrile (400 µL). Samples were vortexed (2,000 rpm, 5 min), clarified by centrifugation (18,000 × g, 8 min), and the supernatants were vacuum dried. The dried samples were reconstituted in 100 µL of 50/50/0.1 water/acetonitrile/formic acid and transferred to new autosampler vials for analysis.

An aliquot of each sample (10 µL) was injected onto a Poroshell 120 SB-C18 analytical column (catalog no. 683775-902, Agilent, 2.1 × 150 mm, 2.7 µm). Analytes were eluted at a flow rate of 400 µL/min using an optimized gradient of solvent A (water, 0.1% formic acid) and solvent B (acetonitrile, 0.1% formic acid; min/%B: 0/10, 2/10, 6/100, 7/10, 15/10) on a 1290 Infinity HPLC system (Agilent Technologies). The column effluent was directed to an OptiFlow Turbo V electrospray ionization source connected to a QTRAP 5500+ triple quadrupole mass spectrometer (AB Sciex LLC) acquiring mass spectra in targeted multiple reaction monitoring (MRM) mode in the positive polarity. Detection of fragmented ions originating from each compound at specific LC retention times was used to ensure specificity and accurate quantification in complex biological samples.

Chromatographic peak areas from the raw mass spectrometry data were integrated and extracted using Analyst (AB Sciex). Standard curves for each compound were generated by plotting the known amount in each standard against the ratio of measured chromatographic peak areas to that of the corresponding internal standard (analyte peak area/IS peak area). The trendline equation was then used to simultaneously normalize to the internal standards and calculate the absolute concentrations of PT-13 in tissue.

### Treatment Study

Female rTg4510 mice were treated with 15 mg/kg PT-13 SC or vehicle injections three days per week, twice daily, from three to five months of age. The study originally began with n = 10 mice per group. Two mice in the PT-13 group died, likely due to external stress. Due to lesion formation in the cervical area upon repeated injections, the dosage regimen was reduced to once daily three days per week when the mice reached six months of age.

### Behavioral Studies

#### Y-Maze Spontaneous Alternation

Vehicle- and PT-13-treated rTg4510 mice were tested to compare spatial memory between the two groups. Mice were acclimated to the Y-maze testing room for one hour prior to the trial. Each mouse was introduced to the long arm and then allowed to explore the maze for five minutes. Video recordings of each trial were scored blinded to provide an unbiased assessment of spontaneous alternation. Spontaneous alternation was defined as three consecutive entries into all three different arms (i.e., ABC, BCA, CAB, etc.). The Y-maze was cleaned thoroughly with 70% ethanol between trials and allowed to dry prior to the next trial (25).

#### Rotarod

Vehicle- and PT-13-treated rTg4510 mice were tested on the rotarod to compare motor performance between the two groups. Mice were acclimated for one hour in the testing room containing the Economex Rotarod (Columbus Instruments, Columbus, OH, USA) with a rotating spindle 7.3 cm in diameter. The rotarod study was conducted over three days. During the first two days, each animal was placed on the rotating spindle for a 30-second acclimation period, followed by 5 minutes walking at a constant speed of 5 rpm (115 cm/min). This process was repeated twice each training day with a 30-minute interval between trials. For the baseline measurement on day three, each animal began at 5 rpm with acceleration at a rate of 0.1 rpm/s (2.3 cm/min/s) until the mouse fell off the rotarod. There was a 30-minute inter-trial interval between the three test trials per mouse. Latency to fall was calculated by excluding the lowest time for each mouse, with the mean of the remaining two trials used as the baseline performance. The rotarod was cleaned thoroughly with 70% ethanol between trials and allowed to dry prior to the next trial (40).

#### Open Field Test

Vehicle- and PT-13-treated rTg4510 mice were tested to compare locomotor abilities between the two groups. Mice were acclimated to the testing room for one hour prior to the trial. The testing room was lit by 90 lux yellow light and was free of non-yellow light sources. Each mouse was placed in the center of an open-field arena consisting of four square chambers, each measuring 27 × 27 cm, arranged in a 2 × 2 grid and constructed from white plastic. A camera was mounted 1.83 m above the floor and centered over the open field. EthoVision XT software captured and analyzed animal movement over a 15-minute recording period for each individual test (41).

#### Morris Water Maze

Spatial learning was assessed using the Morris Water Maze in the campus animal behavioral core. Mice were acclimated to the testing room for one hour prior to the trial. The maze consisted of a circular tank (101 cm in diameter) filled to 10 cm below the rim with 27°C water made opaque by the addition of non-toxic, soluble white paint. A circular escape platform (10 cm in diameter) was submerged 1 cm below the water surface at a fixed location in the southeast quadrant of the tank. Mice were assessed on their ability to locate the platform across five training days, with 6 trials per day and a minimum 30-minute inter-trial interval per mouse. Mice were allowed to swim for 60 seconds to locate and remain on the platform for 3 seconds; if the platform was not located within 60 seconds, the mouse was placed on the platform for 15 seconds before being returned to the home cage. Movement was tracked using EthoVision XT software, and latency to find the platform was recorded manually across all training sessions (41).

#### Probe Trial

Following completion of Morris Water Maze training, a probe trial was conducted. The platform was removed from the southeast quadrant, and each mouse was allowed to swim freely for 60 seconds one hour after the final training trial. Time spent in the target quadrant, the quadrant previously containing the platform, was measured to assess memory of the platform location. After 60 seconds, the mouse was removed from the pool, dried, and returned to its home cage. Movement within each quadrant was tracked using EthoVision XT software (41).

### Euthanasia of Mice

Mice were euthanized with an overdose of Avertin as anesthesia, followed by cervical dislocation for brains used in biochemical assays, or by perfusion with saline for one minute and 4% PFA for four minutes for brains used in histological analyses. Perfused brains were preserved in 4% PFA at 4°C, then transferred to 20% sucrose buffer. Once equilibrated overnight, brains were dried, flash-frozen in 4-methylbutane, and stored at −80°C until sectioning for immunohistochemistry.

### Protein Extraction from Brain Tissue

Cortex tissue was weighed and homogenized in 1× TBS buffer (1:10 w/v) containing protease inhibitor (PI) (Sigma, catalog no. P8340) and phosphatase inhibitor (PhI) (Sigma, catalog no. P0044) using a Dounce glass homogenizer (WHEATON® Tenbroeck Tissue Grinder, 2 mL) on ice for 3 minutes. Samples were then centrifuged at 15,000 × g for 60 minutes at 4°C, and the supernatant was collected as the TBS fraction. The pellets were resuspended in 1× RIPA buffer (Cell Signaling Technology, catalog no. 9806, diluted from 10× with H_2_O) containing PI and PhI, incubated on ice for 40 minutes, and centrifuged at 15,000 × g for 60 minutes at 4°C. The supernatant was collected as the lysis fraction. The remaining pellet was resuspended in 2% SDS (1:5 w/v) and centrifuged at 15,000 × g for 30 minutes at 4°C; the supernatant was collected as the SDS fraction. Protein concentrations were measured using a BCA assay kit, and samples were aliquoted and stored at −80°C until further use.

### Western Blot Analyses

Protein expression was analyzed using protein extracts from the lysis and SDS fractions obtained by the protein extraction protocol described above. Two µL of protein sample was combined with 0.5 µL of 2 M DTT, 2.5 µL of 4× LDS, and 5 µL of H_2_O in a PCR tube. Samples were heated at 90°C for 10 minutes to denature proteins and then separated by gel electrophoresis using Bolt™ Bis-Tris Plus Mini Protein Gels, 4-12% (Thermo Fisher Scientific, Waltham, MA). Proteins were transferred to PVDF membranes using the iBlot 2 NC Regular Stacks (Thermo Fisher Scientific, Waltham, MA) at 20 V for 1 minute, then 23 V for 2 minutes. Membranes were blocked in 5% BSA in 1× TBST for 1 hour at room temperature. After three 5-minute washes with TBST, membranes were incubated overnight at 4°C with the following primary antibodies: TAU-5 (Thermo Fisher, catalog no. AHB0042, 1:500), AT8 (Thermo Fisher, catalog no. MN1020, 1:500), Beta-Actin (Thermo Fisher, catalog no. AM4302, 1:1000), Synaptophysin (Thermo Fisher, catalog no. MA5-14532, 1:1000), PSD-95 (Thermo Fisher, catalog no. MA1-045, 1:1000), and CD68 (Thermo Fisher, catalog no. MA5-13324, 1:1000). After three washes with TBST, membranes were incubated for 1 hour at room temperature with secondary goat anti-rabbit or anti-mouse IgG (Bio-Rad Laboratories, 1:3000) diluted in 5% BSA in TBST. Blots were developed using an ECL system (Bio-Rad Laboratories, Hercules, CA) and visualized using an iBright imaging device (Thermo Fisher, Waltham, MA) (39).

### Immunohistochemistry

Fixed brains were mounted onto a chuck with Optimal Cutting Temperature (OCT) solution and sectioned using a cryostat. Hippocampal slices were placed into netwells within a 12-well plate containing 2 mL of 1× TBS buffer (diluted 1:10 from 10× TBS, pH 7.4) and washed three times for 5 minutes each. Slices were then incubated in a quenching solution consisting of 10% methanol and 3% hydrogen peroxide in 50 mM Tris (pH 7.4) for 10 minutes at room temperature on a nutator, followed by three additional 5-minute washes in 1× TBS. For blocking, tissue sections were incubated in 4% normal goat serum (Vector Labs, catalog no. S-1000-20) in TBS with 0.1% Triton X-100 for 1 hour at room temperature on a nutator. Sections were then incubated with the primary antibody AT8 (Thermo Fisher, catalog no. MN1020, 1:500) diluted in 2% normal goat serum in TBS with 0.1% Triton X-100 for 24 hours at 4°C on a nutator. After three 5-minute washes in 1× TBS, sections were incubated with the secondary antibody, biotin-conjugated goat anti-mouse (Vector Labs, catalog no. PK-6102, 1:4000), diluted in 2% goat serum in TBS with 0.02% Triton X-100 for 2 hours at room temperature on a nutator. Thirty minutes before use, the ABC reagent was prepared from the Vectastain® Elite ABC-HRP Kit (200 µL Buffer A + 200 µL Buffer B in 10 mL 1× TBS) and equilibrated at room temperature. After three 5-minute TBS washes, sections were incubated in the ABC reagent for 1 hour at room temperature on a nutator. Following three additional 5-minute TBS washes, slices were incubated in 3,3’-diaminobenzidine (DAB) solution (10 mg DAB with 50 µL H_2_O_2_ in 50 mL 1× TBS) for 2 minutes at room temperature on a nutator, followed by quenching in 1× TBS. Slices were mounted onto slides and, once dried, dehydrated through a graded ethanol series (70%, 95%, and 100% ethanol, 5 minutes each) followed by xylene (5 minutes). Samples were coverslipped with Permount solution. Images were acquired with a confocal microscope, and AT8 staining was quantified using ImageJ.

### Statistical Analysis

Graphs and statistical analyses were generated using GraphPad Prism software (version 8.0.1). All data are represented as the mean ± standard deviation. Sample sizes for each experiment are indicated in the corresponding figure legends. For all behavioral analyses, data were analyzed by Student’s t-test followed by Tukey’s post hoc test.

## Acknowledgements

We thank Dr. R Kayed for contributing TOMA and TTC-M tau antibodies for our in vitro testing in this study, M. Diamond for providing tau biosensor cell lines, and the W. Cohn and the USC Mann Multi-Omics Mass Spectrometry Core for technical and experimental services contributing to this study. We also thank S. Magaki and C.K. Willimas from the UCLA Pathology Department for contributing AD tissues for these studies. In vitro ADMET experiments were performed by Pharmaron. We are grateful to the Epstein Family Alzheimer’s Disease Research Foundation for financial support for this research through grant awards USC.CA101985.PR0002622 [PMS].

## Author Contributions

NM conducted synthesis, biosensor cell assays, and in vivo work. RF advised and assisted with the in vivo study including the IACUC protocol and guidance in injections and behavior studies. JW generated recombinant tau oligomer. RK assessed behavior study videos blinded to provide a non-biased assessment. AA assisted with proteasome work negative-stain EM. Dr. QLM helped develop the in vivo study and assisted in purifying the cortex tissue from the in vivo study. WC conducted Mass Spectrometry. GP played a major role in the in vivo study design. MJ provided fundamental insight for the in vivo study and trained NM on perfusions and IHC. SKA provided guidance for synthesis and in vitro ADMET characterization. PMS provided funding, overarching guidance, and manuscript writing.

