## Supplementary Figures for "Tau Disaggregation by a CNS-Permeable Small Molecule Reduces Fibril and Oligomer Burden and Preserves Proteostasis and Behavior"

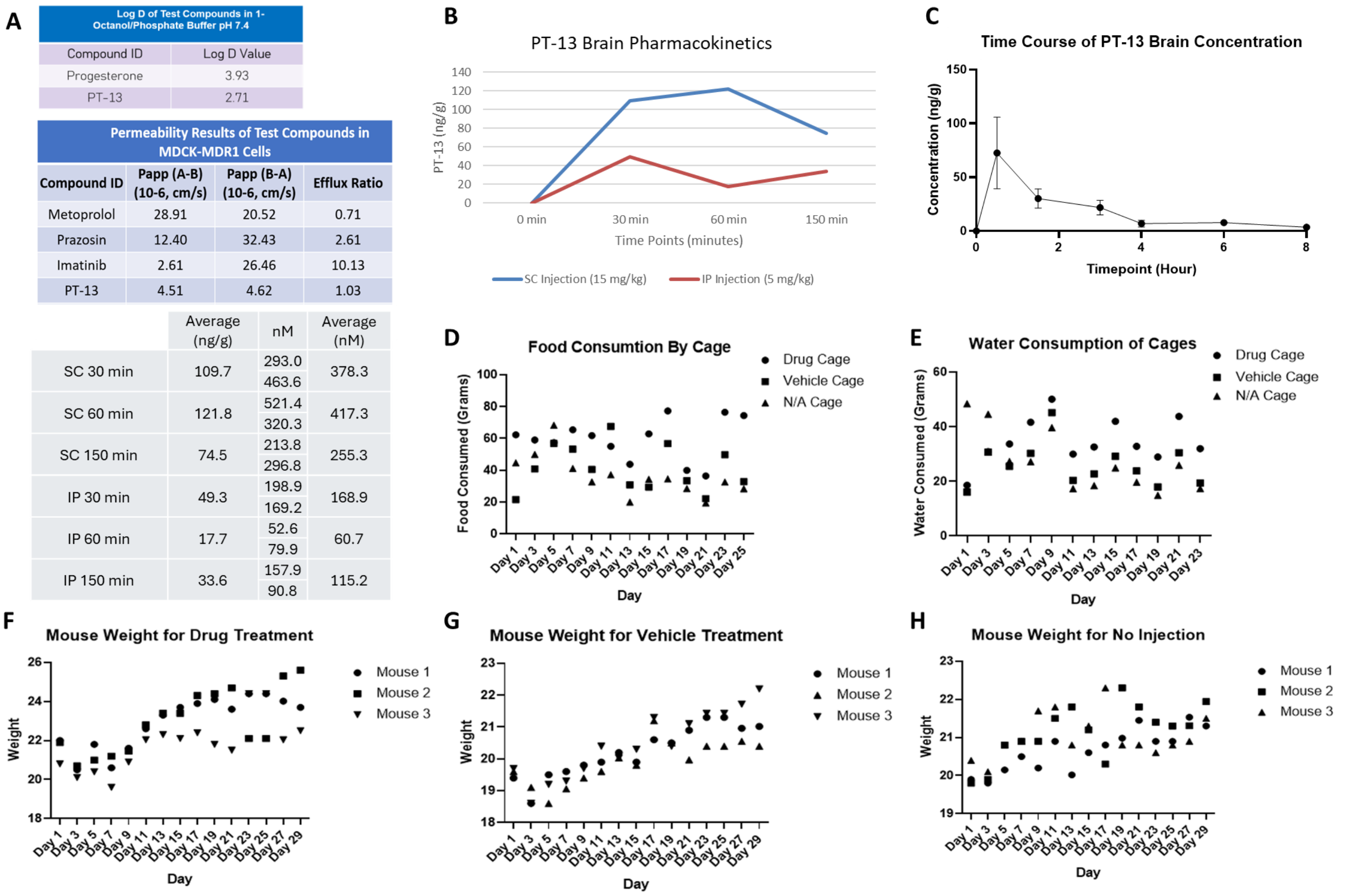

| Log D of Test Compounds in 1-Octanol/Phosphate Buffer pH 7.4 | |
|---|---|
| Compound ID | Log D Value |
| Progesterone | 3.93 |
| PT-13 | 2.71 |

| Permeability Results of Test Compounds in MDCK-MDR1 Cells | | | |
|---|---|---|---|
| Compound ID | Papp (A-B) (10-6, cm/s) | Papp (B-A) (10-6, cm/s) | Efflux Ratio |
| Metoprolol | 28.91 | 20.52 | 0.71 |
| Prazosin | 12.40 | 32.43 | 2.61 |
| Imatinib | 2.61 | 26.46 | 10.13 |
| PT-13 | 4.51 | 4.62 | 1.03 |

| | Average (ng/g) | nM | Average (nM) |
|---|---|---|---|
| SC 30 min | 109.7 | 293.0<br>463.6 | 378.3 |
| SC 60 min | 121.8 | 521.4<br>320.3 | 417.3 |
| SC 150 min | 74.5 | 213.8<br>296.8 | 255.3 |
| IP 30 min | 49.3 | 198.9<br>169.2 | 168.9 |
| IP 60 min | 17.7 | 52.6<br>79.9 | 60.7 |
| IP 150 min | 33.6 | 157.9<br>90.8 | 115.2 |

**Supplementary Figure 1. Pilot data demonstrating PT-13 BBB permeability and dosage tolerability.** (A) In vitro ADMET data. PT-13 demonstrates an efflux ratio of 1.0 in MDCK-MDR1 cells similar to positive control compounds, Metoprolol and Prazosin, indicating effective membrane crossing and stable retention in cells. Log D determination in 1-Octanol/Phosphate Buffer, pH 7.4, indicates excellent cell permeability compared to control molecule, Progesterone. (B) Initial BBB permeability pilot. Mice injected with highest possible dose through subcutaneous injection (SC) or intraperitoneal injection (IP). Injected dose was 5 mg/kg PT-13 by IP and 15 mg/kg SC. PT-13 from brain tissue was quantified by MRM mass spectrometry. C57BL/6J mice n = 2 per timepoint for each injection site. (C) Extended BBB permeability measures for preferred injection route. C57BL/6J mice were injected with a single subcutaneous dose of 15 mg/kg PT-13. Brain concentrations of PT-13 measured at the indicated time points post-injection are shown (n = 3 female mice per time point). (D-H) Dosage tolerance was assessed in PT-13 and control animals by monitoring food and water intake, D and E, in C57BL/6J administered at 15 mg/kg PT-13 was well tolerated over 29 days. No observable signs of toxicity were seen. F-H, mice continued normal weight gain throughout the pilot study for PT-13 and vehicle treatment.

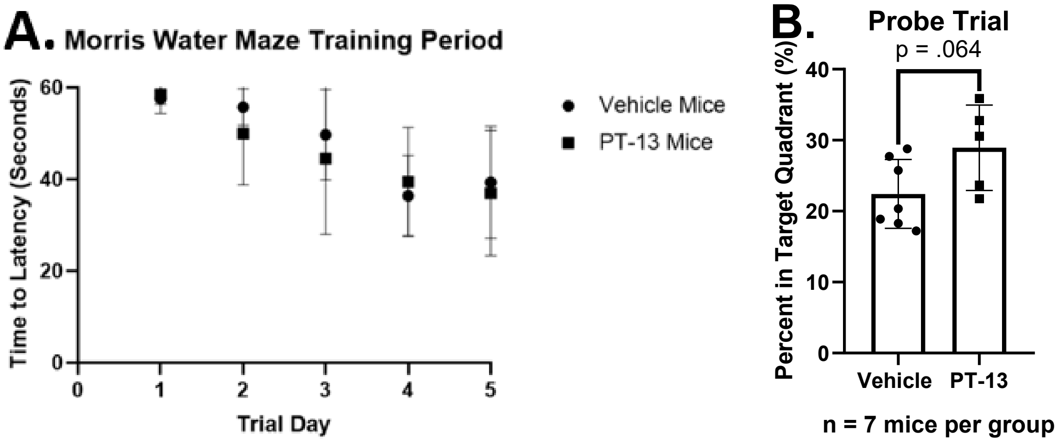

**Supplemental Figure 2:** Drug treatment induces no statistically significant difference in the time of latency for locating the platform in the Morris Water Maze.

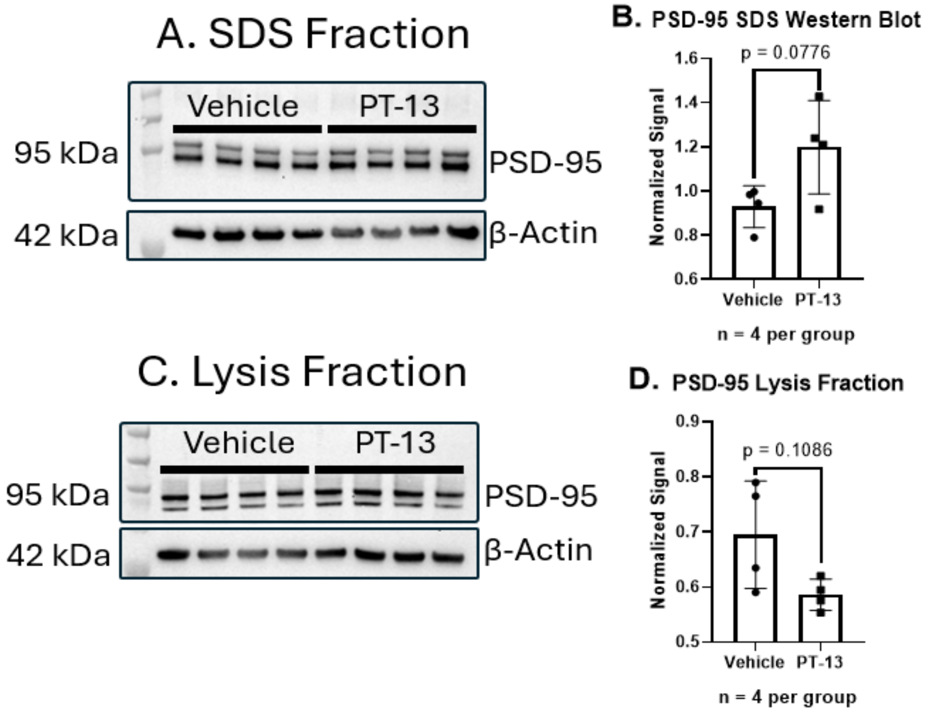

**Supplemental Figure 3:** PT-13 treatment shows a trending shift in PSD-95 concentration between the SDS and Lysis fractions extracted from the cortex tissue of rTg4510 mice. PSD-95 signal was normalized to beta-actin.

**Supplementary Tables**

**Table 1: Biological Activity Data for Compounds M1-M40**

| Compound ID | Structure | Molecular Weight | Biological Data |
|---|---|---|---|
| **M1 'CNS-16'** | 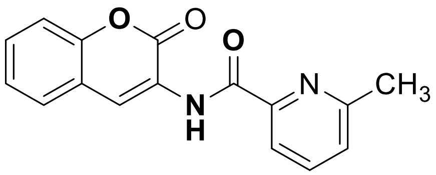 | 280.28 | 20% Inhibition against AD Seeding on average |
| **M2 'PT-31'** | 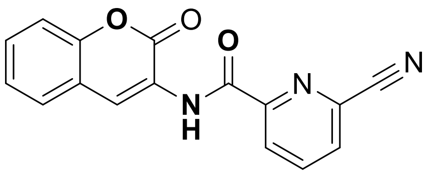 | 291.26 | 60% Inhibition against AD Seeding on average |
| **M3** | 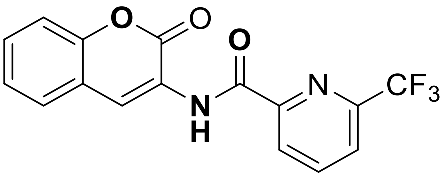 | 334.25 | 30% Increase in AD Seeding on average |
| **M4 'PT-13'** | 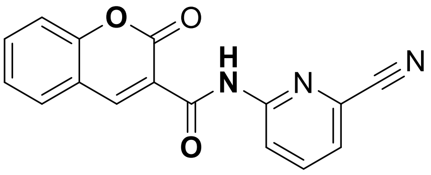 | 291.26 | 85% Inhibition against AD Seeding on average |
| **M5 'PT-92'** | 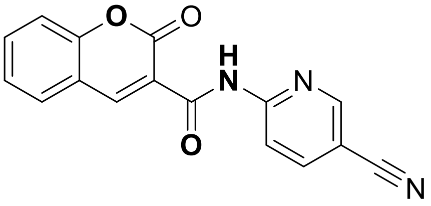 | 291.26 | 50% Inhibition against AD Seeding on average |
| **M6 'PT-97'** | 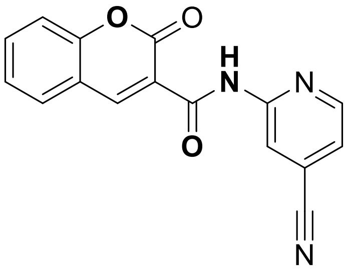 | 291.26 | No notable effect on AD seeding |

| Compound ID | Structure | Molecular Weight | Biological Data |
|---|---|---|---|
| **M7** | 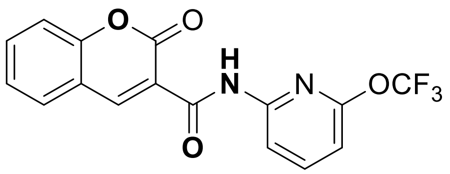 | 350.25 | 25% Inhibition against AD Seeding on average |
| **M8** | 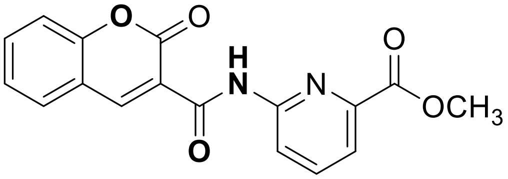 | 324.29 | No notable effect on AD seeding |
| **M9** | 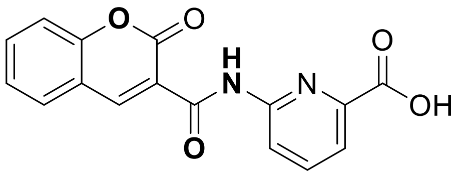 | 310.26 | 30% Inhibition against AD Seeding on average |
| **M10** | 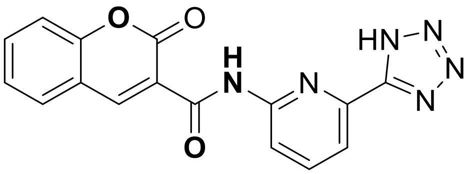 | 334.29 | 40% Inhibition against AD Seeding on average |
| **M11** | 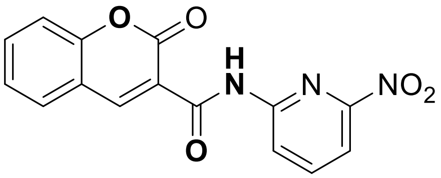 | 311.25 | No notable effect on seeding |
| **M12** | 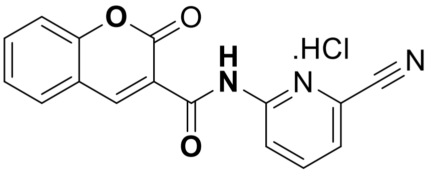 | 327.72 | 85% Inhibition against AD Seeding on average |
| **M13** | 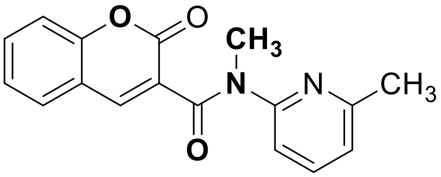 | 294.31 | 10% Increase in AD Seeding on average |

| Compound ID | Structure | Molecular Weight | Biological Data |
|---|---|---|---|
| **M14** | 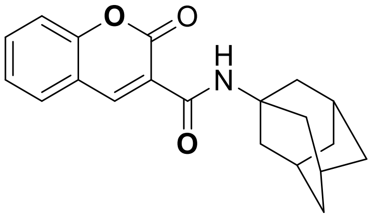 | 323.39 | No notable effect on AD seeding |
| **M15** | 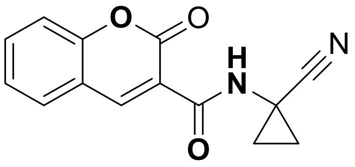 | 254.24 | 35% Inhibition against AD Seeding on average |
| **M16** | 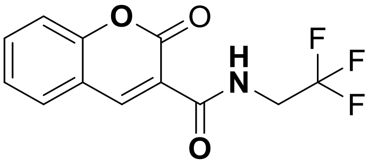 | 271.19 | 35% Inhibition against AD Seeding on average |
| **M17** | 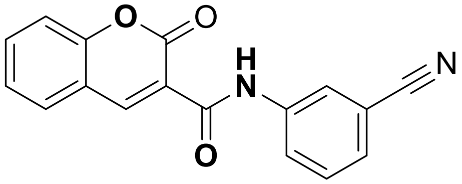 | 290.27 | No notable effect on seeding |
| **M18** | 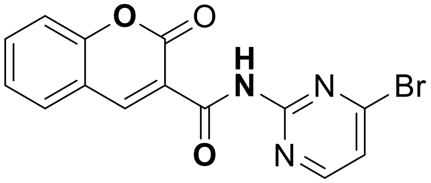 | 346.14 | No notable effect on seeding |
| **M19** | 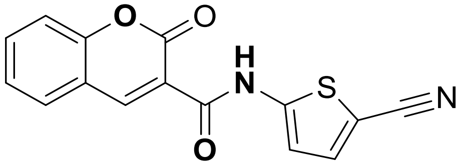 | 296.30 | 40% Inhibition against AD Seeding on average |
| **M20** | 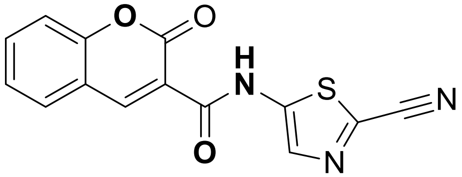 | 297.28 | 50% Inhibition against AD Seeding on average |

| Compound ID | Structure | Molecular Weight | Biological Data |
|---|---|---|---|
| **M21 'PT-46'** | 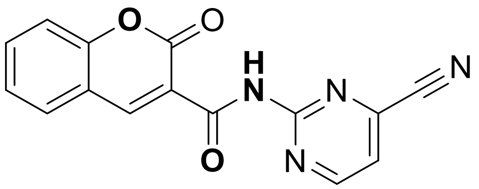 | 292.25 | 20% Inhibition against AD Seeding on average |
| **M22** | 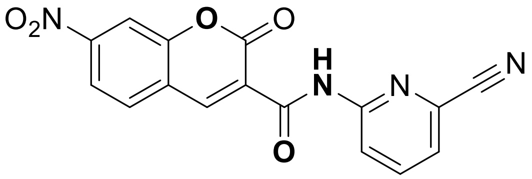 | 336.26 | 30% Increase in AD Seeding on average |
| **M23** | 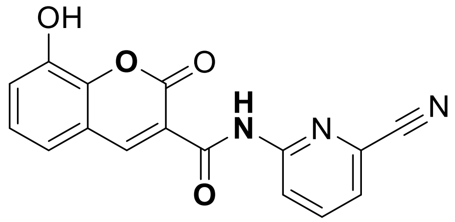 | 307.26 | 50% Inhibition against AD Seeding on average |
| **M24** | 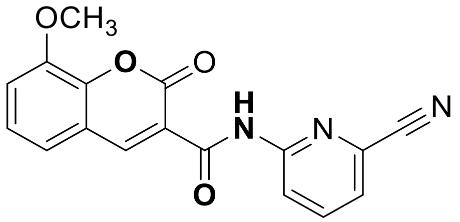 | 321.29 | No notable effect on seeding |
| **M25** | 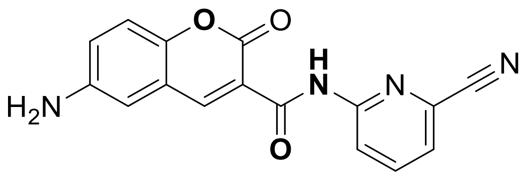 | 306.28 | 50% Inhibition against AD Seeding on average |
| **M26** | 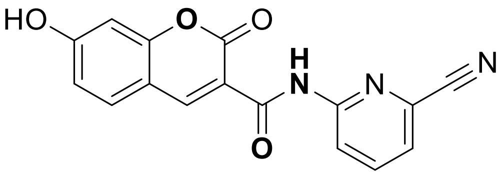 | 307.26 | 55% Inhibition against AD Seeding on average |
| **M27** | 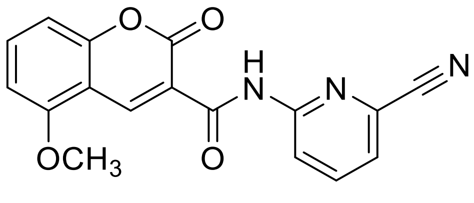 | 321.29 | 40% Inhibition against AD Seeding on average |

| Compound ID | Structure | Molecular Weight | Biological Data |
|---|---|---|---|
| **M28** | 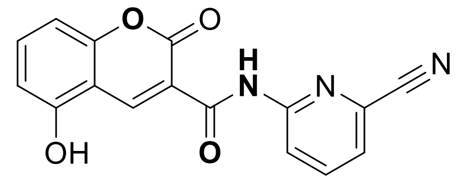 | 307.26 | No notable effect on seeding |
| **M29** | 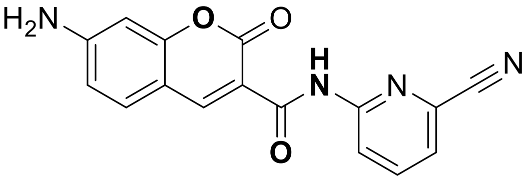 | 306.28 | 50% Increase in AD Seeding on Average |
| **M30** | 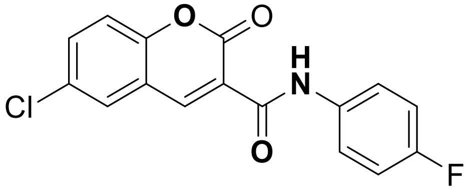 | 317.70 | 20% Increase in AD Seeding on average |
| **M31** | 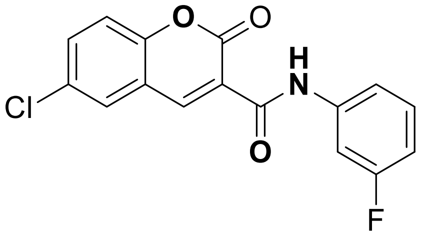 | 317.70 | 30% Inhibition against AD Seeding on average |
| **M32** | 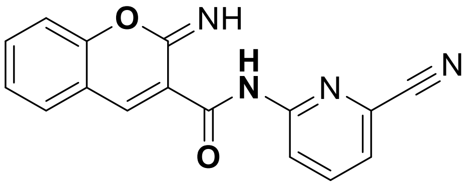 | 290.28 | 50% Inhibition against AD Seeding on average |
| **M33** | 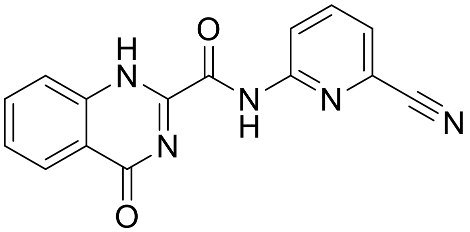 | 291.27 | No notable effect on seeding |
| **M34** | 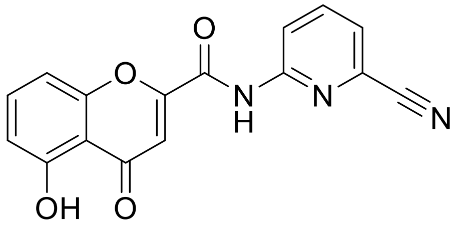 | 291.26 | No notable effect on seeding |

| Compound ID | Structure | Molecular Weight | Biological Data |
|---|---|---|---|
| **M35** | 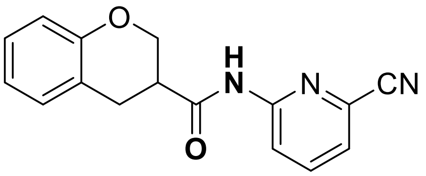 | 279.2990 | No notable effect on seeding |
| **M36** | 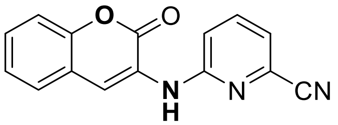 | 263.2560 | No notable effect on seeding |
| **M37**<br>**'PT-243'** | 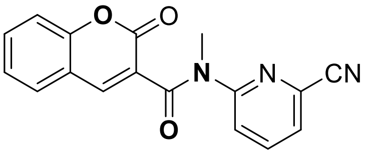 | 305.2930 | No notable effect on seeding |
| **M38** | 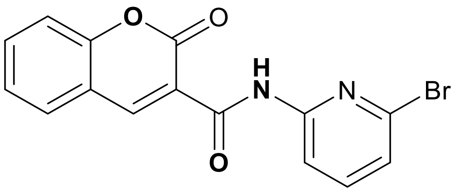 | 345.152 | 30% Increase in AD Seeding on average |
| **M39** | 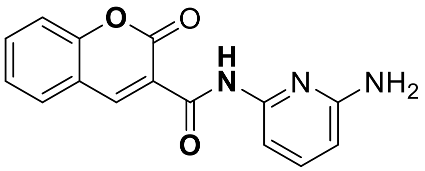 | 281.271 | 40% Inhibition against AD seeding on average |
| **M40** | 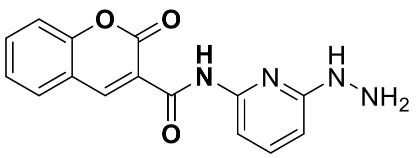 | 296.286 | 70% Inhibition against AD seeding on average |
